# Simple-to-use CRISPR-SpCas9/SaCas9/AsCas12a vector series for genome editing in *Saccharomyces cerevisiae*

**DOI:** 10.1101/2021.05.07.443192

**Authors:** Satoshi Okada, Goro Doi, Shitomi Nakagawa, Emiko Kusumoto, Takashi Ito

## Abstract

Genome editing using the CRISPR/Cas system has been implemented for various organisms and becomes increasingly popular even in the genetically tractable budding yeast *Saccharomyces cerevisiae*. Since each CRISPR/Cas system recognizes only the sequences flanked by its unique protospacer adjacent motif (PAM), a certain single system often fails to target a region of interest due to the lack of PAM, thus necessitating the use of another system with a different PAM. Three CRISPR/Cas systems with distinct PAMs, namely SpCas9, SaCas9, and AsCas12a, have been successfully used in yeast genome editing and their combined use should expand the repertoire of editable targets. However, currently available plasmids for these systems were individually developed under different design principles, thus hampering their seamless use in the practice of genome editing. Here we report a series of Golden Gate Assembly-compatible backbone vectors designed under a unified principle to exploit the three CRISPR/Cas systems in yeast genome editing. We also created a software to assist the design of genome-editing plasmids for individual target sequences using the backbone vectors. Genome editing with these plasmids demonstrated practically sufficient efficiency in both insertion of gene fragments to essential genes and complete deletion of an open reading frame. The backbone vectors with the software would thus provide a versatile toolbox to facilitate the seamless use of SpCas9, SaCas9, and AsCas12a in various types of genome manipulation, especially those that are difficult to perform with conventional techniques in yeast genetics.

## Introduction

Clustered Regularly Interspaced Short Palindromic Repeats (CRISPR) is an adaptive immune system in eubacteria and archaebacteria that functions to counteract foreign nucleic acids, such as those of invading bacteriophages (Jinek *et al*. 2012). The CRISPR array encodes guide RNAs (gRNAs) that form complexes with Cas (CRISPR-associated) proteins. In a number of CRISPR/Cas systems, the Cas–gRNA complexes function as DNA endonuclease to cleave double-stranded DNA. The cleavage target is specified by the sequence of gRNA. Accordingly, co-expression of a Cas protein and its cognate gRNA can introduce a DNA double strand break (DSB) at a specific position in the genome. Cleaving the genome at a specific site is the key process of genome editing. Because of the ease of specifying the target sequence, the CRISPR/Cas systems are being widely used for genome editing in a variety of organisms (Hsu *et al*. 2014).

Protospacer adjacent motif (PAM) is a short DNA sequence that flanks the target sequence defined by the gRNA and is required for the Cas–gRNA complex to recognize its target sequence (Sternberg *et al*. 2014). Nucleotide sequence of PAM is different from one CRISPR/Cas system to another. SpCas9 from *Streptococcus pyogenes* has a G-rich PAM (NGG) that follows the 3′ end of the target sequence (Jinek *et al*. 2012). AsCas12a from *Acidaminococcus sp*. has a T-rich PAM (TTTV) that precedes the 5′ end of the target sequence (Zetsche *et al*. 2015). SaCas9 from *Staphylococcus aureus* has a PAM with an intermediate GC content (NNGRRT) that flanks the 3′ end of the target sequence (Ran *et al*. 2015).

An absolute prerequisite for genome editing to insert a gene fragment to a specific site is a PAM in the vicinity of the target site. It, however, often happens that no appropriate PAM for a single CRISPR/Cas system is found in the region of interest. If multiple systems with different PAMs are available, it would theoretically become much easier to find a PAM and hence the target sequence to introduce a DNA DSB for gene fragment insertion. Indeed, a simple calculation indicates the power of the combined use of SpCas9, SaCas9, and AsCas12a in genome editing of budding yeast *Saccharomyces cerevisiae* (see below).

These three CRISPR/Cas systems have been already implemented for the budding yeast (DiCarlo *et al*. 2013; Laughery *et al*. 2015; Generoso *et al*. 2016; Jessop-Fabre *et al*. 2016; Świat *et al*. 2017; Degreif *et al*. 2018; Verwaal *et al*. 2018). However, the vectors for these systems were independently developed in different laboratories. Consequently, the methods and design principles for genome-editing plasmid construction (e.g., copy number, selection marker, promoter, cloning sites, etc.) are different from one to another. In practice, such differences often hamper seamless, stress-free use of the most suitable system to a given target site of interest. If all the three systems can be used under a unified manner, then the genome-editing processes will be substantially accelerated.

Based on these theoretical and practical needs, we developed in this study a series of four backbone vectors under a unified design principle to seamlessly exploit SpCas9, SaCas9, and enAsCas12a in yeast genome editing. A single highly efficient method, Golden Gate Assembly, is applicable to construct genome-editing plasmids on these backbones. To facilitate the design of synthetic oligodeoxyribonucleotides (ODNs) required for the Golden Gate Assembly process, we developed a simple software that automatically calculates the ODN sequences corresponding to a given target sequence. We demonstrated that genome-editing plasmids thus constructed were efficient enough for routine use in both gene knock-in at essential genes and complete deletion of open reading frames (ORFs).

## Materials and methods

### Yeast strains

Yeast strains used in this study are listed in Table S1. All strains are derived from *Saccharomyces cerevisiae* BY4741 or BY4742 (Brachmann *et al*. 1998). Standard culture media were used in this study (Guthrie and Fink 1991). Conventional gene deletion was performed using a PCR-based method (Longtine *et al*. 1998). Plasmids used for yeast strain construction are listed in Table S2.

### Construction of backbone vectors for genome editing

ODNs used in this study are listed in Table S3. All ODNs for plasmid construction were purchased from Sigma-Aldrich Japan (Tokyo, Japan) and Eurofins Genomics K. K. (Tokyo, Japan). The four backbone vectors for genome editing were constructed using the seamless cloning with HiFi DNA Assembly (E2621, New England Biolabs, Ipswich, MA, USA). Restriction enzymes used for plasmid construction were purchased from New England Biolabs. PCR fragments used for plasmid construction were amplified by Q5 DNA polymerase (M0491, New England Biolabs) according to manufacturer’s instruction. *Escherichia coli* competent cells NEB 5-alpha (C2987, New England Biolabs), NEB Stable (C3040, New England Biolabs), or Champion DH5α high (CC5202, SMOBIO Technology, Hsinchu City, Taiwan) were used for transformation to amplify and extract plasmids. Plasmids were extracted by FastGene Plasmid Mini Kit (FG-90502, Nippon Genetics, Tokyo, Japan). Plasmids used in this study are listed in Table S2. The DNA sequence files of the backbone vectors for genome editing are available on our repository at GitHub (https://github.com/poccopen/Genome_editing_plasmid_for_budding_yeast).

### Selection of target sequences for genome editing

For the insertion of mNeonGreen-encoding sequence into the *CSE4* gene, we selected target sequences from the region encoding the unstructured N-terminal loop of Cse4 protein (Zhou *et al*. 2011; Yan *et al*. 2019). For the insertion of mScarlet-I-encoding sequence into the *CDC3* gene, we first performed secondary structure prediction by JPred4 (Drozdetskiy *et al*. 2015) of Cdc3 protein and then selected target sequences from the region encoding the N-terminal region with no predicted secondary structure.

For designing single guide RNAs (sgRNAs) for SpCas9 and SaCas9, CRISPRdirect (Naito *et al*. 2015) was used to select target sequences. For designing CRISPR RNAs (crRNAs) for enAsCas12a, CRISPOR (Concordet and Haeussler 2018) was used to select target sequences. Target sequences for genome editing used in this study are listed in Table S4.

### Construction of genome-editing plasmids

All genome-editing plasmids were constructed using the seamless cloning with Golden Gate Assembly using NEB Golden Gate Assembly Kit (BsaI-HF v2) (E1601, New England Biolabs). The ODNs for Golden Gate Assembly were automatically designed with an in-house software.

### Yeast transformation for genome editing

Yeast transformation was carried out as described previously (Gietz and Woods 2002) with slight modifications. Yeast cells were cultured overnight in 2 mL of YPAD liquid medium (10 g/L Bacto Yeast Extract, #212750, Thermo Fisher Scientific, Waltham, MA, USA; 20 g/L Bacto Peptone, #211677, Thermo Fisher Scientific;100 mg/L adenine sulfate, #01990-94, Nacalai tesque, Kyoto, Japan; and 20 g/L glucose, Nacalai tesque) at 25°C with shaking at 250 rpm. The 2-mL overnight culture was centrifugated and the supernatant was removed. The cell pellet was resuspended in 0.5 mL of 0.1 M lithium acetate solution (#127-01545, FUJIFILM Wako Chemicals, Osaka, Japan). The cell suspension was incubated at 30°C for 30 min. Fifty microliters of cell suspension were thoroughly mixed with 50 μL of 1 M lithium acetate, 50 μL of 1 M dithiothreitol (#14128-04, Nacalai tesque), 5 μL of Yeastmaker Carrier DNA (10 mg/mL, #630440, Takara Bio, Kusatsu, Japan), 1 μL of genome-editing plasmid (200–600 ng), 45 μL of PCR-generated donor fragment for gene fragment insertion (1–10 μg, typically 5 μg) or ORF deletion (2.5 μg), and 300 μL of polyethylene glycol 4000 (#11574-15, Nacalai tesque). PCR fragments were amplified by Q5 DNA polymerase (New England Biolabs) or KOD One (KMM-101, TOYOBO, Osaka, Japan) according to manufacturer’s instructions. The samples were incubated at 30°C for 45 min followed by a 15-min incubation at 42°C. After centrifugation and removal of supernatant, the cell pellets were resuspended with 50 μL of SC−Ura medium without carbon source (7.4 g/L Yeast nitrogen base without amino acids, #291940, Thermo Fisher Scientific; 855 mg/L CSM−Ura powder, DCS0161, FORMEDIUM, Hunstanton, UK; and 111 mg/L adenine sulfate, Nacalai tesque) and spread on a SCGal−Ura agar plate (20 g/L galactose, #075-00035, FUJIFILM Wako Chemicals; 6.7 g/L Yeast nitrogen base without amino acids, Thermo Fisher Scientific; 770 mg/L CSM−Ura powder, FORMEDIUM; 100 mg/L adenine sulfate, Nacalai tesque; and 20 g/L agar, #010-08725, FUJIFILM Wako Chemicals). The plates were incubated at 30°C for 4 days. The colonies were picked and streaked as patches on SCGal−Ura agar plates, and then incubated at 30°C for 1–2 days followed by colony PCR to check successful genome editing. Colony PCR was performed using Q5 DNA polymerase (New England Biolabs) or KOD One (TOYOBO) according to manufacturer’s instructions. The PCR-positive clones were cultured overnight in 2 mL of YPAD liquid medium. An aliquot (10 μL) of the overnight culture was spotted and streaked on a YPAD agar plate for single colony isolation (30°C for 2 days). Single colonies were picked and streaked on YPAD agar plate and SCDex−Ura agar plate (20 g/L glucose, #16806-25, Nacalai tesque; 6.7 g/L Yeast nitrogen base without amino acids, Thermo Fisher Scientific; 770 mg/L CSM−Ura powder, FORMEDIUM; 100 mg/L adenine sulfate, Nacalai tesque; and 20 g/L agar, #010-08725, FUJIFILM Wako Chemicals) to check the loss of the genome-editing plasmid. The Ura^−^ clones were re-examined by colony PCR to be successfully genome-edited. The colony PCR-positive Ura^−^ clones were used in the subsequent experiments.

### Plasmid extraction from yeast cells

Plasmids were extracted from yeast cells by Easy Yeast Plasmid Isolation Kit (#630467, Takara Bio) and transformed into *E. coli* competent cells (Champion DH5α high, SMOBIO Technology).

### Fluorescence microscopy and image processing

Image acquisitions of yeast cells were performed on a microscope (Ti-E, Nikon, Tokyo, Japan) with a 100× objective lens (CFI Apo TIRF 100XC Oil, MRD01991, Nikon), a sCMOS camera (ORCA-Fusion BT, C15440-20UP, Hamamatsu photonics, Hamamatsu, Japan), and a solid-state illumination light source (SOLA SE II, Lumencor, Beaverton, OR, USA). Image acquisition was controlled by NIS-Elements version 5.3 (Nikon). The binning mode of the camera was set at 2×2 (0.13 μm/pixel). Z-stacks were 13×0.3 μm. For imaging of Cse4-mNeonGreen, a filter set (LED-YFP-A, Semrock, Rochester, NY, USA) was used with excitation light power set at 20% and exposure time set at 200 msec/frame. For imaging of Cdc3-mScarlet-I, a filter set (LED-TRITC-A, Semrock) was used with excitation light power set at 7% and exposure time set at 70 msec/frame. For DIC (differential interference contrast) image acquisition, exposure time was set at 20 msec/frame. DIC images were captured only at the middle position of the Z-stacks.

Image processing and analysis were performed using Fiji (Schindelin *et al*. 2012). To generate 2-dimensional images of fluorescence channel from Z-stacks, background subtraction (sliding paraboloid radius set at 10 pixels with disabled smoothing) and maximum projection using 13 Z-slices were performed. Maximum projected fluorescence images and corresponding smoothed DIC images were superimposed. After global adjusting of brightness and contrast and cropping of the images, sequences of representative images were generated.

### Editable fraction of yeast genome with three CRISPR/Cas systems

*S. cerevisiae* reference genome sequence available at *Saccharomyces* genome database (SGD) (S288C strain, version R64-2-1, http://sgd-archive.yeastgenome.org/sequence/S288C_reference/genome_releases/S288C_reference_genome_R64-2-1_20150113.tgz) without mitochondrial genome and plasmid sequences were searched for PAMs (NGG for SpCas9, NNGRRT for SaCas9, and TTTV for AsCas12a). Both strands were included in the PAM search. After identification of the PAM sequence, nucleotides in a defined distance from the PAM were assigned as candidate nucleotides for editing. In genome editing, it is critical for a successfully-edited target sequence in the genome not to be cleaved again by the Cas–gRNA complex bearing the gRNA corresponding to the original, unedited target sequence. We thus defined a nucleotide as a candidate for editing if its substitutions leading to mismatches with the gRNA can significantly reduce the efficiency of re-cleavage by the Cas–gRNA complex. For SpCas9 and SaCas9, the nucleotides at one to eleven nt away from the PAM and the nucleotides consisting the PAM were defined as the candidates based on previous reports (Anderson *et al*. 2015; Zheng *et al*. 2017; Tycko *et al*. 2018) (Figures S1A and S1B). For AsCas12a, the nucleotides that are one to seventeen nt away from the PAM and the nucleotides consisting the PAM were defined as the candidates (Kim *et al*. 2016; Kleinstiver *et al*. 2016; Bin Moon *et al*. 2018) (Figure S1C). Degenerate nucleotides in the PAMs (i.e., N, R, and V) were excluded from the calculation (Figures S1A, S1B, and S1C). The total number of the candidate nucleotides for editing is summarized in Table S5 and Figure S1D.

### ORFs editable at their 5′ ends

*S. cerevisiae* ORF sequence collection available at SGD (http://sgd-archive.yeastgenome.org/sequence/S288C_reference/orf_dna/orf_genomic_all.fasta.gz) was used for the search of ORFs that can be edited at their 5′ ends. ORFs on the mitochondrial genome and ORFs on the two-micron plasmids were omitted from the analysis. The total number of ORFs analyzed in this study was 6,881. Candidate nucleotides for editing were searched in each sequence (ORF and upstream 1,000 and downstream 1,000 nt) by the method described above. When at least one nucleotide within the start codon ‘ATG’ was assigned as the candidate for editing, the ORF was categorized as an ORF that is editable at its 5′ end. The total number of the ORFs editable at the 5′ ends is summarized in Table S6 and Figure S1E.

### Data availability

The four backbone vectors are available from NBRP Yeast Resource Center (https://yeast.nig.ac.jp/yeast/). The source codes of programs for ODN design, PAM search, and 5′-editable ORF search are available from our repository at GitHub (https://github.com/poccopen/Genome_editing_plasmid_for_budding_yeast). Other strains and plasmids are available upon request. The authors state that all data necessary for confirming the conclusions presented here are represented fully within the article.

## Results

### Expansion of editable ORFs by combining three CRISPR/Cas systems

We evaluated the potentials of three well-established CRISPR/Cas systems with distinct PAMs, namely SpCas9 from *Streptococcus pyogenes* (Jinek *et al*. 2012), SaCas9 from *Staphylococcus aureus* (Ran *et al*. 2015), and enhanced AsCas12a (enAsCas12a) derived from *Acidaminococcus sp*. (Kleinstiver *et al*. 2019), in editing the budding yeast genome. For each system, we calculated the number of “editable” nucleotides and estimated the fraction of ORFs amenable to genome editing at their 5′ ends (e.g., N-terminal fusion with a tag protein) (see Materials and Methods). This simple simulation indicated that while SpCas9, SaCas9, and AsCas12a can target the 5′ ends of 4,277 (62.2%), 2,804 (40.7%), and 4,101 (59.6%) of 6,881 ORFs, respectively, their combined use can cover as much as 6,116 ORFs (88.9%) (Figure S1, Table S6). Based on this simulation, we decided to develop a backbone vector series sharing a single design principle to enable the seamless use of the three CRISPR/Cas systems to expand the repertoire of editable genes.

### Design of backbone vectors for yeast genome editing

In the design of the vector series, we defined the following three requirements: 1) both Cas protein and sgRNA/crRNA are encoded on a single plasmid, 2) expression of Cas protein and/or sgRNA/crRNA can be artificially induced, and 3) target sequence of sgRNA/crRNA can be incorporated using the Golden Gate Assembly (Engler *et al*. 2008). Fulfilling these three requirements, we developed four backbone centromeric plasmid vectors marked with *URA3* for the three CRISPR/Cas systems (Figure 1). Note that enAsCas12a was used instead of AsCas12a because of its improved activity at lower temperature suitable to grow budding yeast cells (Kleinstiver *et al*. 2019).

**Figure 1.**
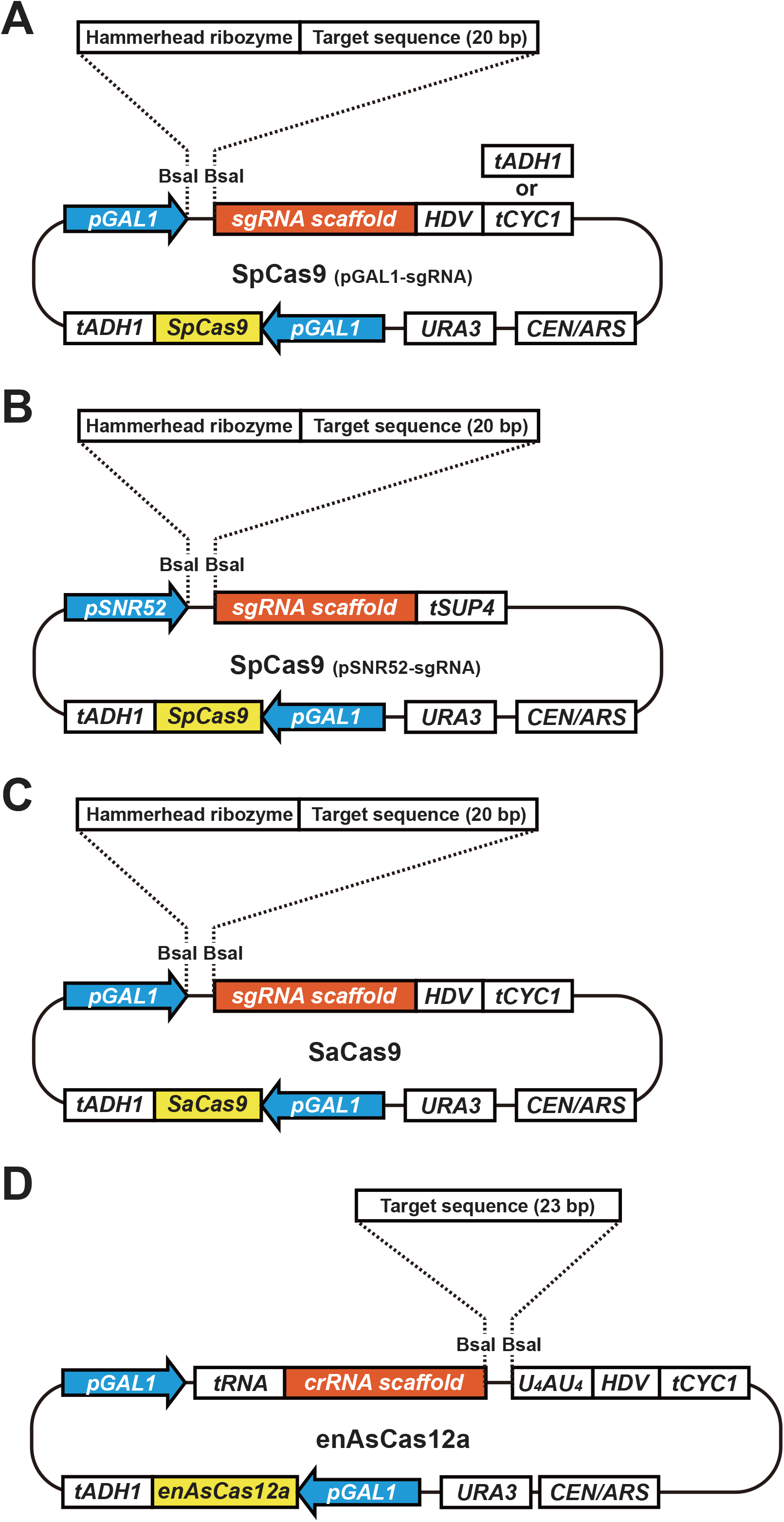
Schematic representation of backbone vectors for genome editing. (A) SpCas9 + pGAL1-sgRNA system. (B) SpCas9 + pSNR52-sgRNA system. (C) SaCas9 system. (D) enAsCas12a system. *pGAL1, GAL1* promoter; *pSNR52, SNR52* promoter; *tADH1, ADH1* terminator; *tCYC1, CYC1* terminator; *tSUP4, SUP4* terminator; HDV, HDV ribozyme; *U*_*4*_*AU*_*4*_, 9-mer encoding “UUUUAUUUU” for improvement of genome-editing efficiency; *URA3, URA3* marker cassette; *CEN/ARS*, centromere and autonomously replicating sequence

As an inducible promoter, we use the well-characterized *GAL1* promoter because it is actively repressed by glucose and strongly activated by galactose in the absence of glucose. Cas-encoding genes on the four vectors are placed under the control of the *GAL1* promoter. Similarly, sgRNA/crRNA precursors on three vectors are controlled by the *GAL1* promoter.

The Golden Gate Assembly uses type IIS restriction enzymes such as BsaI and BbsI (Engler *et al*. 2008). Our vector series harbors two BsaI recognition sites for Golden Gate Assembly. Since the target sequence lies at the 5′ terminal side of the sgRNA scaffold for SpCas9 and SaCas9, one BsaI site is placed just downstream of the *GAL1* or *SNR52* promoter and the other site is placed just upstream of the sgRNA scaffold sequence (Figures 1A, 1B, and 1C). In the case of enAsCas12a, the target sequence is located at the 3′ terminal side of the crRNA scaffold. Accordingly, one BsaI site is placed just downstream of the crRNA scaffold and the other site is placed further downstream (Figure 1D).

Extra sequences at the 5′ and 3′ ends of sgRNA often compromise the efficiency of genome editing and hence should be adequately trimmed. To remove the 5′ extra sequence, a hammerhead ribozyme is inserted at the beginning of sgRNA-containing transcript from the three Cas9 vectors (Figures 1A, 1B, and 1C). To remove the 3′ extra sequence in the transcripts driven by *GAL1* promoter, the hepatitis delta virus (HDV) ribozyme is inserted to the 3′ side of the sgRNA scaffold (Figures 1A and 1C). For the other vector using *SNR52* promoter driven by RNA polymerase III, *tSUP4* is used to define the 3′ end of transcript (Figure 1B). In the enAsCas12a vector, the crRNA is preceded and followed by tRNA(Gly) (Zhang *et al*. 2019) and HDV, respectively, for the removal of extra sequences (Figure 1D). Furthermore, a sequence encoding “UUUUAUUUU” is inserted between the second BsaI site (i.e., 3′ end of the crRNA) and the HDV ribozyme, because this 9-mer sequence was demonstrated to increase the efficiency of genome editing (Bin Moon *et al*. 2018).

### Software to design ODNs for Golden Gate Assembly

To construct a genome-editing plasmid, a pair of ODNs corresponding to its target sequence must be synthesized so that they include 4-nt sequences compatible with the backbone vectors. Furthermore, a hammerhead ribozyme compatible with each target sequence must be designed and included in the ODNs (Figure 2A). To facilitate this complicated process without the risk of human errors, we created a simple software that automatically calculates the ODNs for a given target sequence (Figure 2B). Upon entering a target sequence with its name followed by the selection of a backbone vector, the software readily provides ODN sequences to be synthesized for the Golden Gate Assembly of a genome-editing plasmid on the selected backbone vector. The software is available from our repository on GitHub (https://github.com/poccopen/Genome_editing_plasmid_for_budding_yeast).

**Figure 2.**
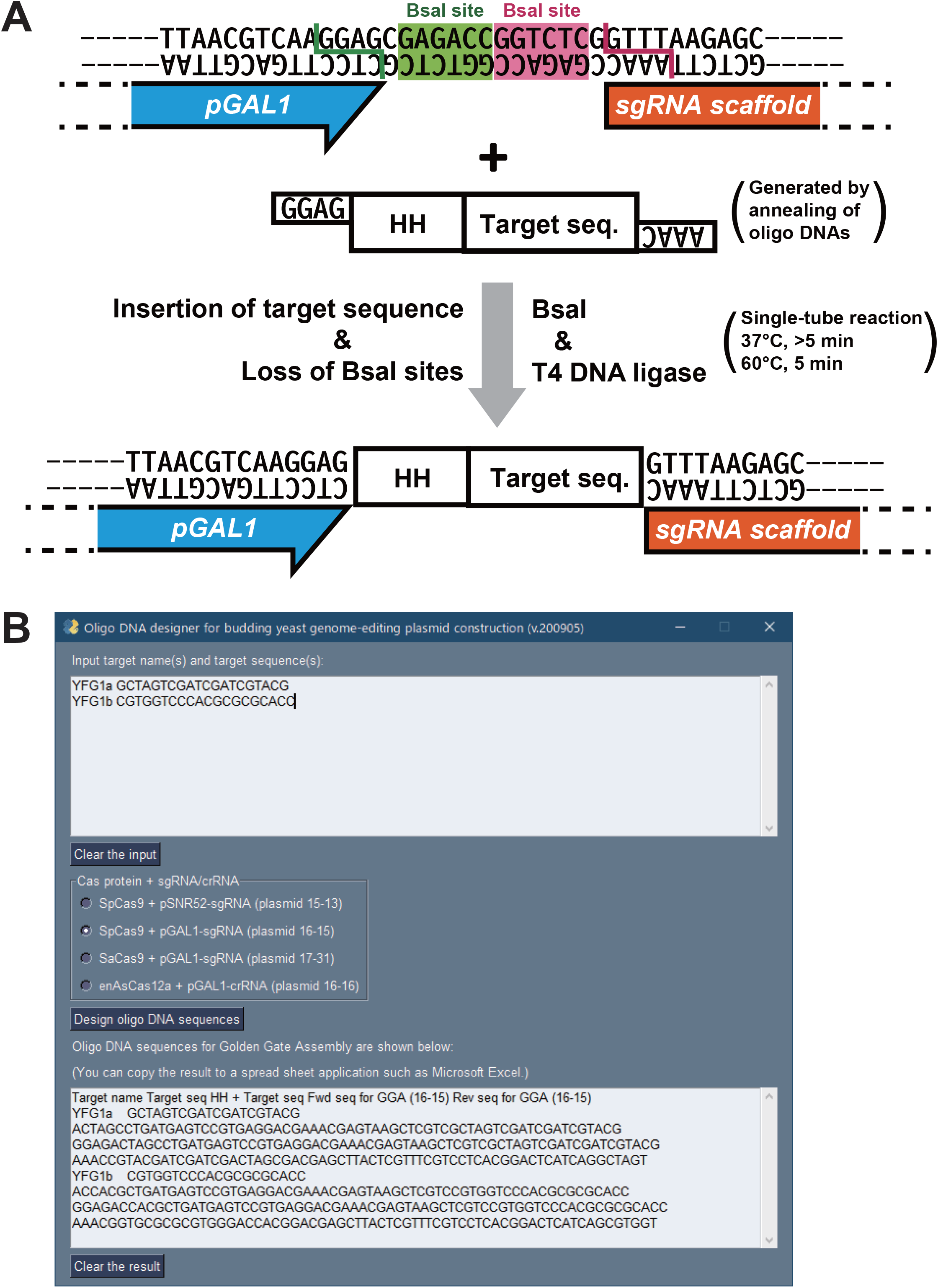
Golden Gate Assembly of genome-editing plasmids and software to design ODN sequences. (A) Process to construct a genome-editing plasmid by Golden Gate Assembly. This panel shows an example for SpCas9 + pGAL1-sgRNA system. HH, hammerhead ribozyme-encoding sequence. (B) Screenshot of the software to automatically design the ODN sequences for Golden Gate Assembly. Upon inputting target sequences with their names to the box at the top followed by the selection of a backbone vector at the middle part, ODN sequences are displayed in the box at the bottom.

### Application example 1: gene insertion by SpCas9 + pGAL1-sgRNA system

Insertion of a DNA fragment to an essential gene at a location other than its ends is difficult to perform even in the budding yeast. As an application example of SpCas9 + pGAL1-sgRNA system, we attempted to insert a fluorescent protein gene into an internal portion of an essential gene. We chose the *CSE4* gene as our target of fluorescent protein gene insertion. The *CSE4* gene encodes a centromere-specific histone H3 variant Cse4 (Stoler *et al*. 1995). It was reported that when a fluorescent protein is fused at the C-terminus of Cse4, the cells show temperature sensitivity (Wisniewski *et al*. 2014). In contrast, when the fluorescent protein is inserted into the unstructured N-terminal loop of Cse4 (Zhou *et al*. 2011; Yan *et al*. 2019), the cells grow normally at higher temperature (Wisniewski *et al*. 2014).

We attempted to insert a gene fragment encoding mNeonGreen, a bright yellow-green fluorescent protein (Shaner *et al*. 2013), into the N-terminal loop of Cse4. We designed 8 target sites in the genic region encoding the N-terminal loop of Cse4 (Figure 3A). To insert the mNeonGreen gene fragment to these sites, we prepared donor PCR fragments harboring 45-bp homology arms at both termini (Figure 3B). For each of the 8 target sites, the yeast cells were co-transformed with the corresponding genome-editing plasmid and donor PCR fragment. The transformants formed a mixture of large and small colonies (Figure S2A). For the target sequence CSE4-1, small colonies showed a significantly higher insertion efficiency (87.5%, n = 24, or 8 colonies in each of 3 biological replicates) than large colonies (4.2%, n = 24, or 8 colonies in each of 3 biological replicates) (Figure S2B).

**Figure 3.**
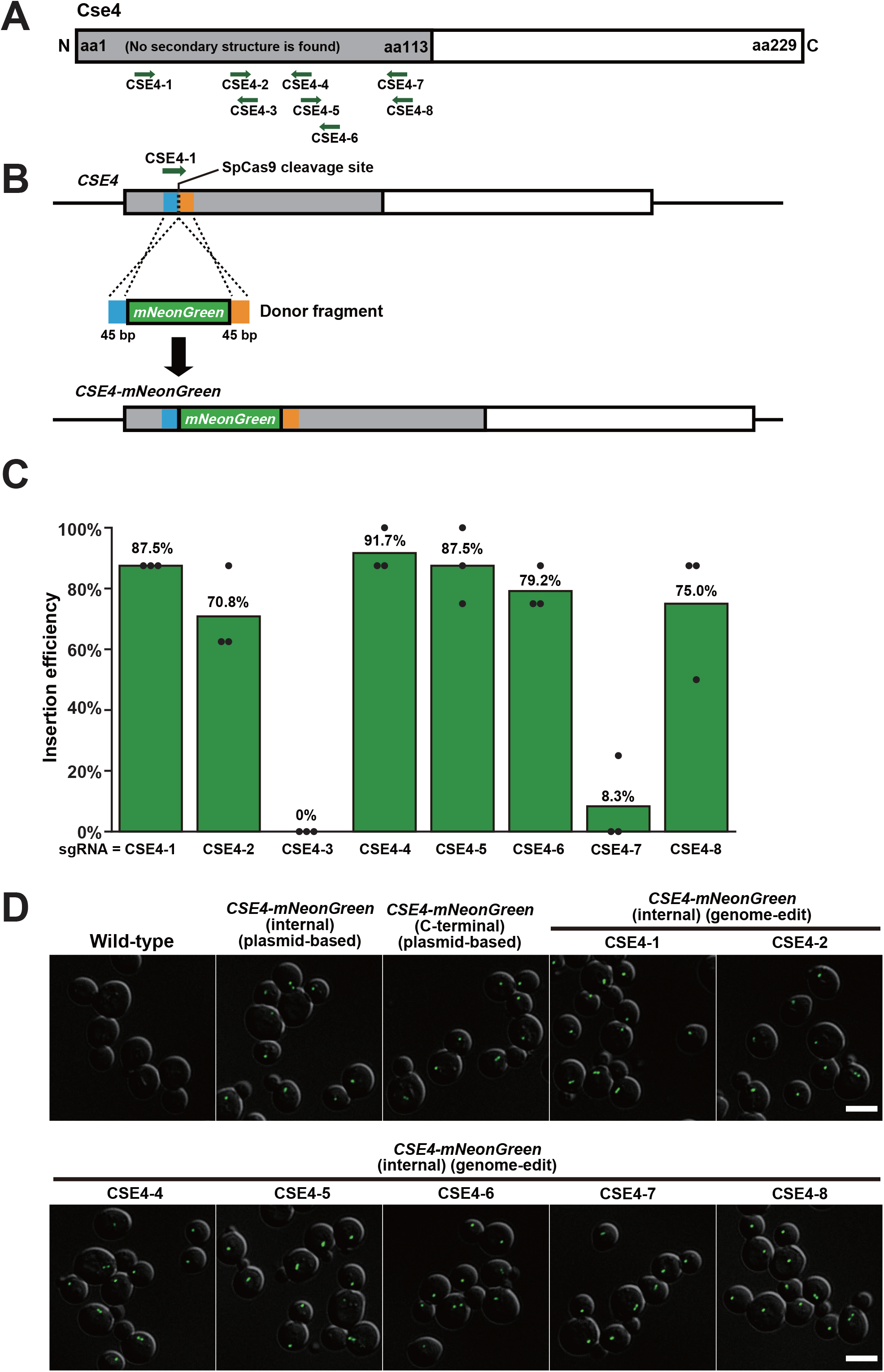
Gene fragment insertion into the essential gene *CSE4* using SpCas9 + pGAL1-sgRNA system. (A) Structure of Cse4 protein. The N-terminal unstructured region is colored in gray. The green arrows represent the positions of the target sequences for genome editing. Leftward and rightward arrows indicate target sequences on the sense and antisense DNA strands, respectively. (B) Gene fragment insertion process by genome editing using SpCas9 at the target sequence CSE4-1 (schematic not proportional to actual size). The mNeonGreen gene fragment to be inserted is colored in green. The 45-bp regions used as the 5′-and 3′-homology arms are colored in blue and orange, respectively. (C) Insertion efficiency at each target sequence. Green bars indicate the average insertion efficiency over three experiments (n = 24 in total). Black dots show the insertion efficiency of each experiment (n = 8 for each). (D) Representative images of the wild-type cells and the *CSE4-mNeonGreen* cells. Images are composed by superimposition of DIC images (gray scale) and mNeonGreen fluorescent images (green). The target sequence names are shown above the images. Scale bar, 5 μm.

Observing the heterogeneity in colony size, we hypothesized that loss of genome-editing function results in loss of cell cycle arrest induced by DSB and its repair and leads to the formation of the large colonies. It was likely that intramolecular recombination between the two *GAL1* promoters led to loss of SpCas9 expression cassette (Figure S2C). To test this hypothesis, we analyzed the structure of the genome-editing plasmids in the cells forming large colonies by restriction enzyme digestion and PCR. All the 24 plasmids derived from large colonies showed the structural change consistent with the predicted deletion caused by recombination between the two *GAL1* promoters (Figures S2D and S2E). Based on these results, we picked only small colonies in the subsequent genome-editing experiments with plasmids harboring two *GAL1* promoters.

Using CSE4-1 as a model target sequence, we also investigated the relationship between the length of homology arms and the insertion efficiency. We used PCR fragments harboring 4 different homology arm lengths (15-, 25-, 35-, and 45-bp) for genome editing. There was a positive correlation between homology arm length and insertion efficiency (Figure S3). A similar positive correlation has been reported between homology arm length and genome-editing efficiency in fission yeast (Hayashi and Tanaka 2019). When a donor PCR fragment harboring 15-bp homology arms was used, no gene insertion was observed (Figure S3). Based on these results, PCR fragments harboring 45-bp homology arms were used in the subsequent genome-editing experiments for gene insertion.

Among the 8 target sequences (Figure 3A, Table S4), the gene insertion efficiency varied from 0% to 91.7% (n = 24 for each target sequence) (Figure 3C, Table S4). Clones with successful gene insertions were obtained for 7 out of the 8 target sequences.

We investigated the phenotype of the successfully genome-edited cells. In all the genome-edited clones tested, the mNeonGreen fluorescence signal was localized as a single spot or a pair of spots in each cell (Figures 3D and S4A). The localization pattern of the mNeonGreen signal was indistinguishable between the genome-edited cells and the cells generated using the conventional method to harbor mNeonGreen at the Cse4 N-terminal loop (Figure S4A). These two genome-modified cells showed comparable growth at 37°C with the wild-type cells, whereas those harboring mNeonGreen at the C-terminus of Cse4 did not (Figure S4B).

### Application example 2: gene insertion by SpCas9 + pSNR52-sgRNA system

As an example of the use of SpCas9 + pSNR52-sgRNA system, we attempted to insert a fluorescent protein-encoding gene into an internal part of an essential gene. We chose the *CDC3* gene as the target of fluorescent protein gene insertion. The *CDC3* gene encodes one of the septin proteins, which form a ring structure along the bud neck (Caviston *et al*. 2003). It was shown that when a fluorescent protein is fused to the C-terminus of Cdc3, the localization of Cdc3 protein becomes abnormal, leading to a morphological defect (Huh *et al*. 2003; Dubreuil *et al*. 2019). In contrast, when a fluorescent protein is inserted into an N-terminal loop of Cdc3, the tagged Cdc3 protein correctly localizes at the bud neck and the cells grow normally without showing any morphological defect (Caviston *et al*. 2003).

We thus attempted to insert a gene fragment encoding a bright red fluorescent protein, mScarlet-I (Bindels *et al*. 2016), into an N-terminal region predicted to lack any secondary structure. We designed 4 target sequences in the genic region corresponding to the N terminal region of Cdc3 (Figure 4A). To insert the mScarlet-I gene fragment, we prepared donor PCR fragments harboring 45-bp homology arms at both termini (Figure 4B). The transformation of yeast cells with a genome-editing plasmid and a corresponding donor PCR fragment resulted in a mixture of large and small colonies (Figure S5A). However, for all the 4 target sequences, there was no statistically significant difference in insertion efficiency between the small and large colonies (Figure S5B).

**Figure 4.**
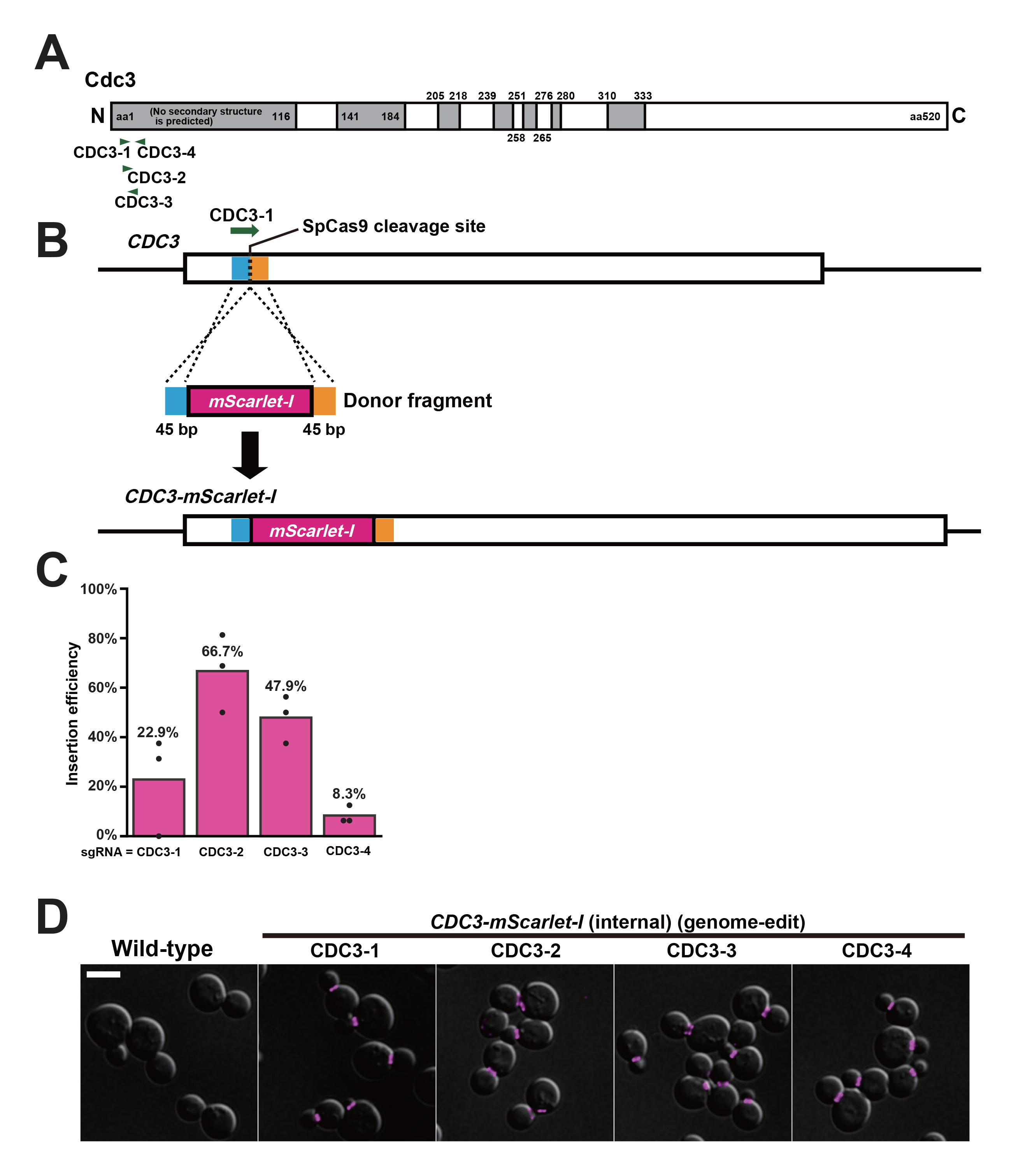
Gene fragment insertion into the essential gene *CDC3* using SpCas9 + pSNR52-sgRNA system. (A) Structure of Cdc3 protein. Regions with no predicted secondary structure are colored in gray. The green arrowheads indicate the positions of target sequences for genome editing. (B) Gene fragment insertion process by genome editing using SpCas9 at the target sequence CDC3-1 (schematic not proportional to actual size). The 45-bp regions used as the 5′-and 3′-homology arms are colored in blue and orange, respectively. (C) Insertion efficiency at each target sequence. Magenta bars indicate the average insertion efficiency over three experiments (n = 48 in total). Black dots show the insertion efficiency of each experiment (n = 16 for each). (D) Representative images of the wild-type cells and the genome-edited *CDC3-mScarlet-I* cells. Images are composed by superimposition of DIC images (gray scale) and mScarlet-I fluorescent images (magenta). The target sequence names are shown above the images. Scale bar, 5 μm.

The gene insertion efficiency varied from 8.3% to 66.7% among the 4 target sequences (n = 48 for each target sequence) (Figure 4C). In all the genome-edited clones examined, mScarlet-I fluorescent signal was localized as a ring structure at the bud neck (Figures 4D and S5C). None of the 12 genome-edited clones (3 clones for each target sequence) exhibited morphological defect (Figure S5C). All the genome-edited clones (4 clones for each target sequence) showed comparable growth at 37°C with the wild-type cells (Figure S5D).

### Application example 3: gene insertion by enAsCas12a system

As an application example of enAsCas12a + pGAL1-crRNA system, we attempted to insert a gene fragment encoding mNeonGreen to the genic regions encoding the N-terminal loop of Cse4, as we did above (Figure 3). We designed 8 target sequences (Figure 5A) and prepared donor PCR fragments harboring 45-bp homology arms at both termini (Figure 5B). The gene insertion efficiency varied from 0% to 87.5% among the 8 target sequences (n = 24 for each target sequence) (Figure 5C). Clones with successful gene insertions were obtained for 6 out of the 8 target sequences (Figure 5C). In all genome-edited clones, the mNeonGreen fluorescence signal was localized as a single spot or a pair of spots in each cell (Figures 5D and S6A) and the growth at 37°C were comparable to the wild-type cells (Figure S6B). All these results were consistent with those obtained for the cells generated using the conventional approach and the SpCas9 + pGAL1-sgRNA system.

**Figure 5.**
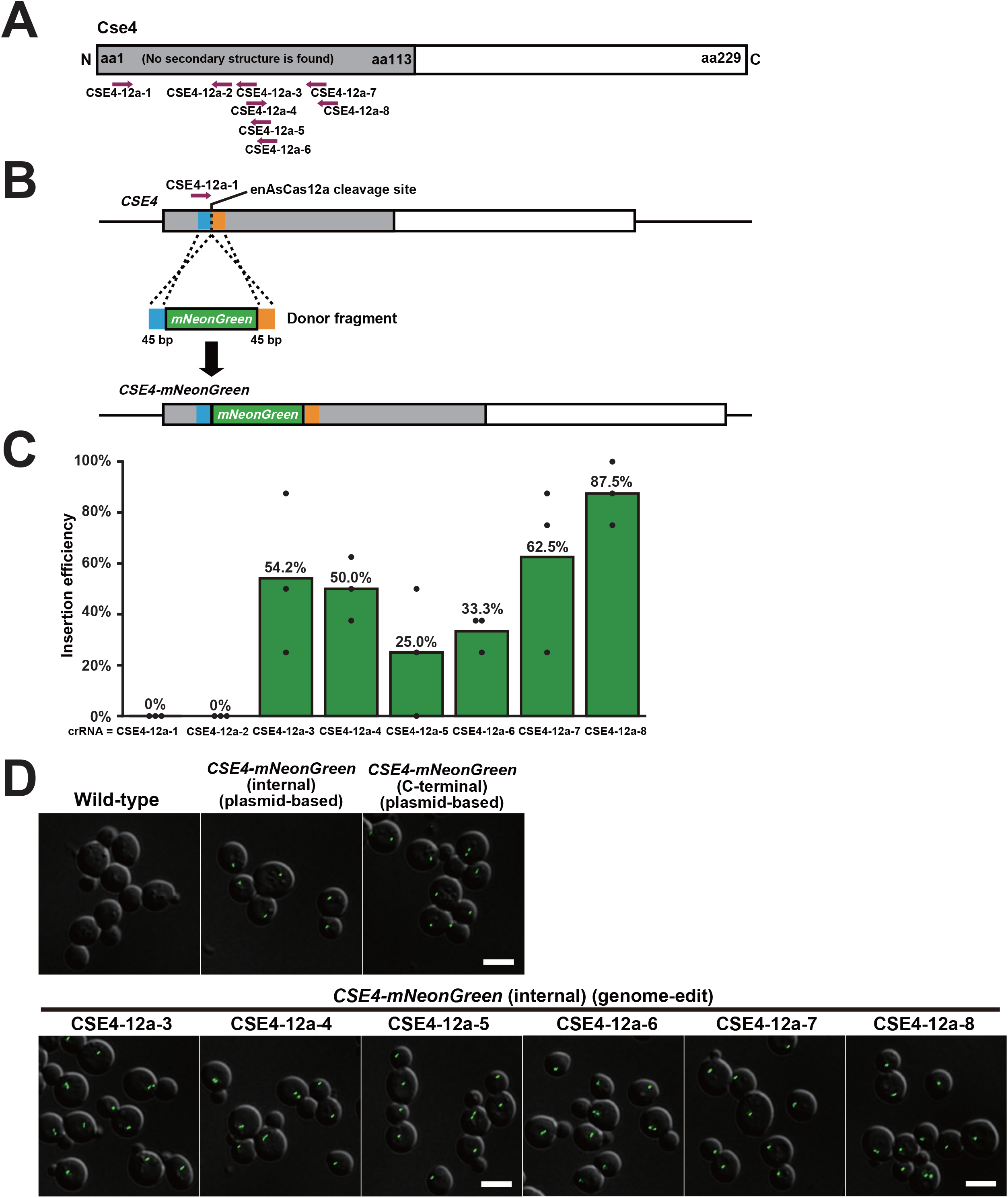
Gene fragment insertion into the essential gene *CSE4* using enAsCas12a system. (A) Structure of Cse4 protein. The N-terminal unstructured region is colored in gray. The dark magenta arrows represent the positions of the target sequences for genome editing using enAsCas12a. (B) Gene fragment insertion process by genome editing using enAsCas12a at the target sequence CSE4-12a-1. The mNeonGreen gene fragment to be inserted is colored in green. The 45-bp regions used as the 5′-and 3′-homology arms are colored in blue and orange, respectively. (C) Insertion efficiency at each target sequence. Green bars indicate the average insertion efficiency over three experiments (n = 24 in total). Black dots show the insertion efficiency of each experiment (n = 8 for each). (D) Representative images of the wild-type cells and the *CSE4-mNeonGreen* cells. Images are composed by superimposition of DIC images (gray scale) and mNeonGreen fluorescent images (green). The target sequence names are shown above the images. Scale bar, 5 μm.

### Application example 4: complete ORF deletion by SaCas9 system

As an example of the use of SaCas9 + pGAL1-sgRNA system, we attempted to delete an entire ORF. We chose the *ADE3* gene as a target of complete ORF deletion. The *ADE3* gene encodes C1-tetrahydrofolate synthase, an enzyme required for adenine biosynthesis (McKenzie and Jones 1977). The cells lacking the *ADE2* gene encoding phosphoribosylaminoimidazole carboxylase, another enzyme in the adenine biosynthesis pathway, form red colonies by accumulating intermediate metabolites of red color (Hieter *et al*. 1985). When the *ADE3* gene is deleted in the cells lacking *ADE2*, colony color returns to white because of the loss of accumulation of the red metabolites (Koshland *et al*. 1985). We attempted to delete the *ADE3* ORF in an *ade2*Δ strain and convert colony color from red to white (Figure 6A). We selected 4 target sequences in the ORF, constructed the corresponding genome-editing plasmids, and used them to transform the *ade2*Δ cells with or without a 100-bp donor PCR fragment composed of the 5′-and 3′-flanking sequences of the ORF (Figure 6A). The transformation with the genome-editing plasmids resulted in the formation of white colonies on galactose-containing plates (Figure 6B, top row). Transformation with a control plasmid YCplac33 failed to form white colonies (Figure 6B). Even when the genome-editing plasmids were used, white colonies did not appear among the transformants on glucose-containing plates (Figure 6B, bottom row). These results indicated that the galactose-inducible SaCas9 system worked as we expected.

**Figure 6.**
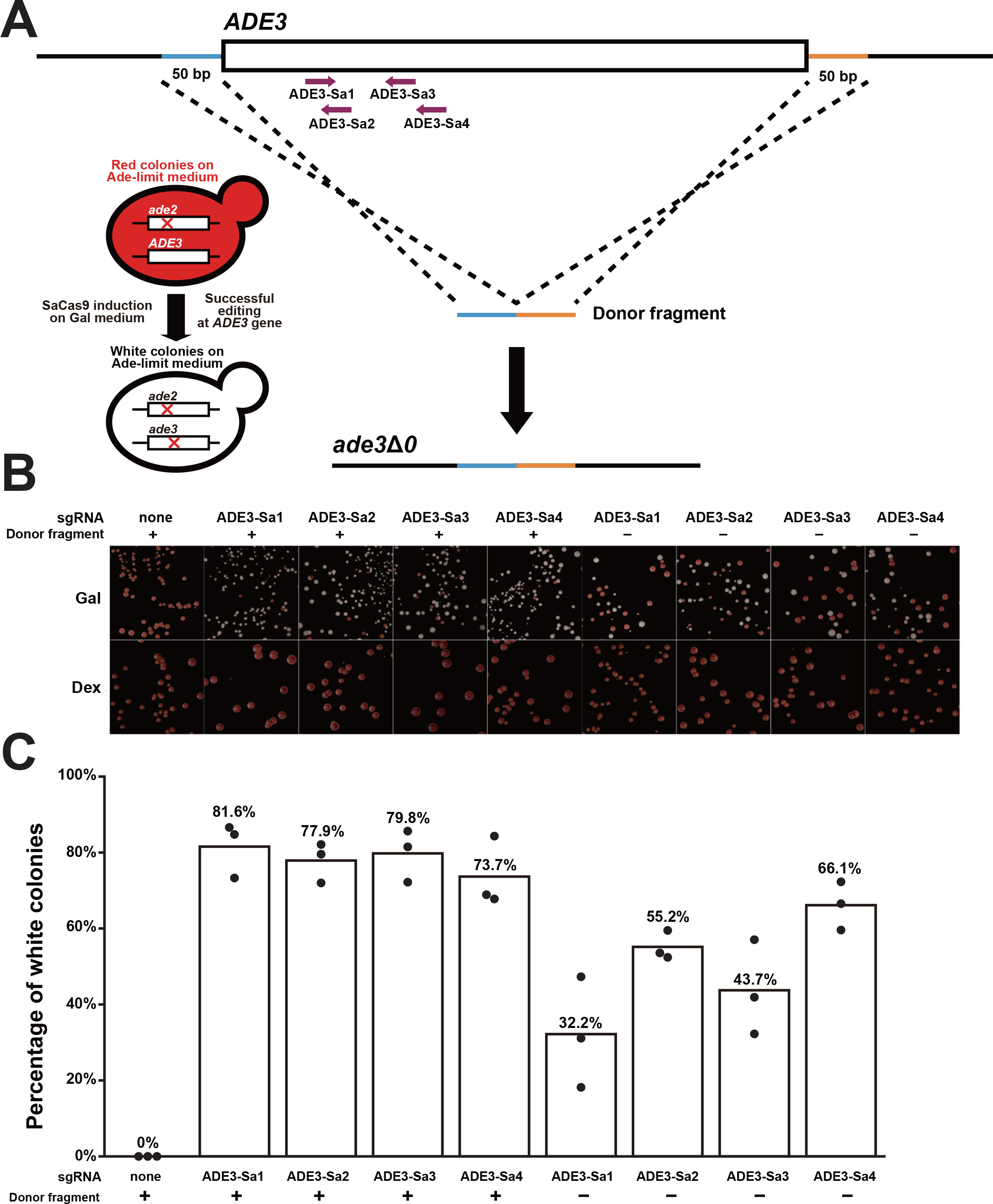
Complete deletion of the entire *ADE3* ORF using SaCas9 system. (A) Process for deleting the entire *ADE3* ORF and converting colony color. The dark magenta arrows represent the positions of the target sequences for SaCas9. The 50-bp regions used as the 5′-and 3′-homology arms of the 100-bp donor PCR fragment are colored in blue and orange, respectively (schematic not proportional to actual size). While *ade2*Δ cells accumulate red pigments on adenine-limited medium, *ade2*Δ*ade3*Δ cells fail to do so and form white colonies. (B) Representative images of colonies after transformation of the genome-editing plasmids with or without the donor PCR products on adenine-limited galactose-containing medium (top, Gal) or on adenine-limited glucose-containing medium (bottom, Dex). The target sequences used for genome editing are shown above the panels. (C) Efficiency of colony color conversion. Each white bar indicates the percentage of white colonies in each experimental condition shown at the bottom. The value is the average of 3 biological replicates indicated by black dots.

When the cells were transformed with a genome-editing plasmid and the donor PCR fragment for the ORF deletion, the proportion of white colonies on galactose-containing plates was in the range of 73.7% to 81.6% (Figure 6C). The formation of white colonies does not necessarily indicate complete deletion of *ADE3* ORF, as small insertion or deletion (indel) could also result in the loss of Ade3 function. To distinguish complete deletion of *ADE3* ORF from small indels, we performed a PCR assay (Figure S7A). PCR products consistent with complete deletion of *ADE3* ORF were obtained in all the 32 white colonies examined (8 colonies for each target sequence) (Figure S7B, top). We also checked the sequence of these PCR products and confirmed the complete loss of *ADE3* ORF (Figure S8).

We also attempted to knock out the *ADE3* gene through non-homologous end joining. For this purpose, we transformed the yeast cells solely with the genome-editing plasmids. In this case, loss of the *ADE3* gene function should be attributable to frameshift mutations caused by indels in the vicinity of SaCas9 cleavage site. The proportion of white colonies on galactose-containing plates was in a range from 32.2% to 66.1%, which is lower than that of the cells co-transformed with the genome-editing plasmids and the donor PCR fragment (73.7%–81.6%) (Figure 6C). We performed a PCR assay to exclude the possibility of large deletions (Figures S7A and S7B, bottom). Sequencing of these PCR products confirmed the presence of small indels (1–2 bp) in the vicinity of the expected SaCas9 cleavage sites (Figure S9).

## Discussion

Here we reported a series of vectors for yeast genome editing using three different CRISPR/Cas systems, namely SpCas9, SaCas9, and enAsCas12a (Figure 1). Since the three systems have distinct PAMs, their combined use expands the repertoire of editable genes, as indicated by our simulation (Figure S1). To facilitate the seamless use of these systems, we constructed a vector series under a unified design principle. First, all the vectors harbor *URA3* marker and *GAL1* promoter, thus sharing the media required for their use. Accordingly, if a single certain system fails to edit a region of interest, one can readily switch to another system without preparing any additional medium. Second, all the vectors are compatible with the highly efficient Golden Gate Assembly, thus making the construction step virtually free from failure. Furthermore, a dedicated software is developed to design ODNs for Golden Gate Assembly of individual genome-editing plasmids on these backbone vectors. Target search with CRISPRdirect (Naito *et al*. 2015) and CRISPOR (Concordet and Haeussler 2018) followed by ODN design with this software would thus streamline the entire process to design genome-editing plasmids.

The realistic schedule of genome editing described in this study is summarized in Table 1. The entire process from designing a genome-editing plasmid to obtaining genome-edited strains can be completed within 2 weeks. This period is substantially shorter than the one required for the traditional yeast genetics approach, especially, in the case of inserting a gene fragment to an essential gene. For instance, when using the own promoter of an essential gene, the traditional approach includes the construction of a cover plasmid carrying the wild-type allele of the essential gene, transformation of the cover plasmid, disruption of the genomic copy of the essential gene, introduction of an adequately-modified allele, and the curing of the cover plasmid, thus taking at least 17 days or, more realistically, >20 days.

**Table 1.** Schedule of genome editing

| Day | Procedures |
| --- | --- |
| 1 | Select target sequences; design and order ODNs |
| 2 |  |
| 3 | Receive ODNs; perform Golden Gate Assembly; transform *E. coli* cells |
| 4 | Inoculate *E. coli* clones for plasmid extraction; inoculate yeast cells for genome-editing transformation |
| 5 | Extract plasmid from *E. coli* clones; prepare donor fragments with PCR; transform yeast cells |
| 6 |  |
| 7 |  |
| 8 |  |
| 9 | Pick up yeast colonies |
| 10 | Perform yeast colony PCR; inoculate yeast cells in YPAD |
| 11 | Spread yeast cells for single colony isolation |
| 12 |  |
| 13 | Pick up single yeast colonies |
| 14 | Perform yeast colony PCR |

It is intriguing to note that the period for genome editing can be further shortened with these plasmids. In this study, we used galactose to strongly activate *GAL1* promoter at the expense of substantially compromised growth compared to that in the presence of glucose, the ideal carbon source for *S. cerevisiae*. Notably, the artificial transcription factor GEV (i.e., a fusion protein composed of Gal4 DNA-binding domain, estrogen receptor, and VP16) can activate the *GAL1* promoter upon estradiol addition in glucose media (Hickman *et al*. 2011). We thus expect that a GEV-bearing strain reconciles efficient induction and rapid growth, thereby further shortening the period required for genome editing using these vectors.

We examined the performance of genome-editing plasmids using these backbone vectors in our attempts to insert gene fragments to essential genes (Figures 3–5) and complete deletion of an ORF (Figure 6). In the case of gene fragment insertion, successfully genome-edited cells were obtained for 17 out of 20 target sequences examined (Figures 3C, 4C, and 5C) and the insertion efficiency exceeded 50% for 11 target sequences. In the case of complete ORF deletion, the efficiency was larger than 70% for all the 4 target sequences tested (Figure 6C). We also examined the growth of genome-edited clones at 30°C and 37°C (Figures S4B, S5D, and S6B). None of the 55 clones showed temperature-sensitive growth, suggesting minimal off-target effects leading to growth defects for the target sequences used in this study. These results proved the practical utility of the vector series developed in this study. We should refer to a practical rule of thumb for successful genome editing using the three backbone vectors bearing two *GAL1* promoters. When using these vectors, intramolecular recombination between the two promoters tends to lead to the formation of large colonies with low efficiency of genome editing (Figure S2). We thus recommend the users of these vectors to simply discard large colonies and select small ones for further analyses because the latter showed significantly higher genome-editing efficiency than the former (Figure S2). While the single *GAL1* promoter plasmid also led to a heterogeneity in colony size, no difference in genome-editing efficiency was observed between large and small colonies (Figure S5).

Our application examples included the insertion of fluorescent proteins into such positions that are neither N-nor C-end of the essential proteins Cse4 and Cdc3 (Figures 3–5). Tagging at inappropriate sites of these proteins was reported to induce temperature-sensitive growth and/or morphological defects. To avoid adverse effects of inserting a fluorescent protein on the recipient protein folding, we took a strategy to select an insertion site from regions demonstrated or predicted to have no secondary structure. All the proteins thus fluorescently-tagged, including those using previously unvalidated sites, showed physiological localization, and the cells thus modified exhibited neither temperature-sensitive growth nor abnormal morphology. These results suggest the general utility of our strategy.

Taken together, the backbone vectors and the software developed in this study would provide a versatile toolbox to facilitate various types of genome manipulation in *S. cerevisiae*, including those difficult to perform with conventional techniques in yeast genetics.

## Acknowledgments

We thank Seiya Kamino, Yuki Hisano, and Masahiro Sakemi for their assistance in plasmid construction and pilot use of the genome-editing systems during the initial phase of the project.

## Supplemental figure legends

**Figure S1.**
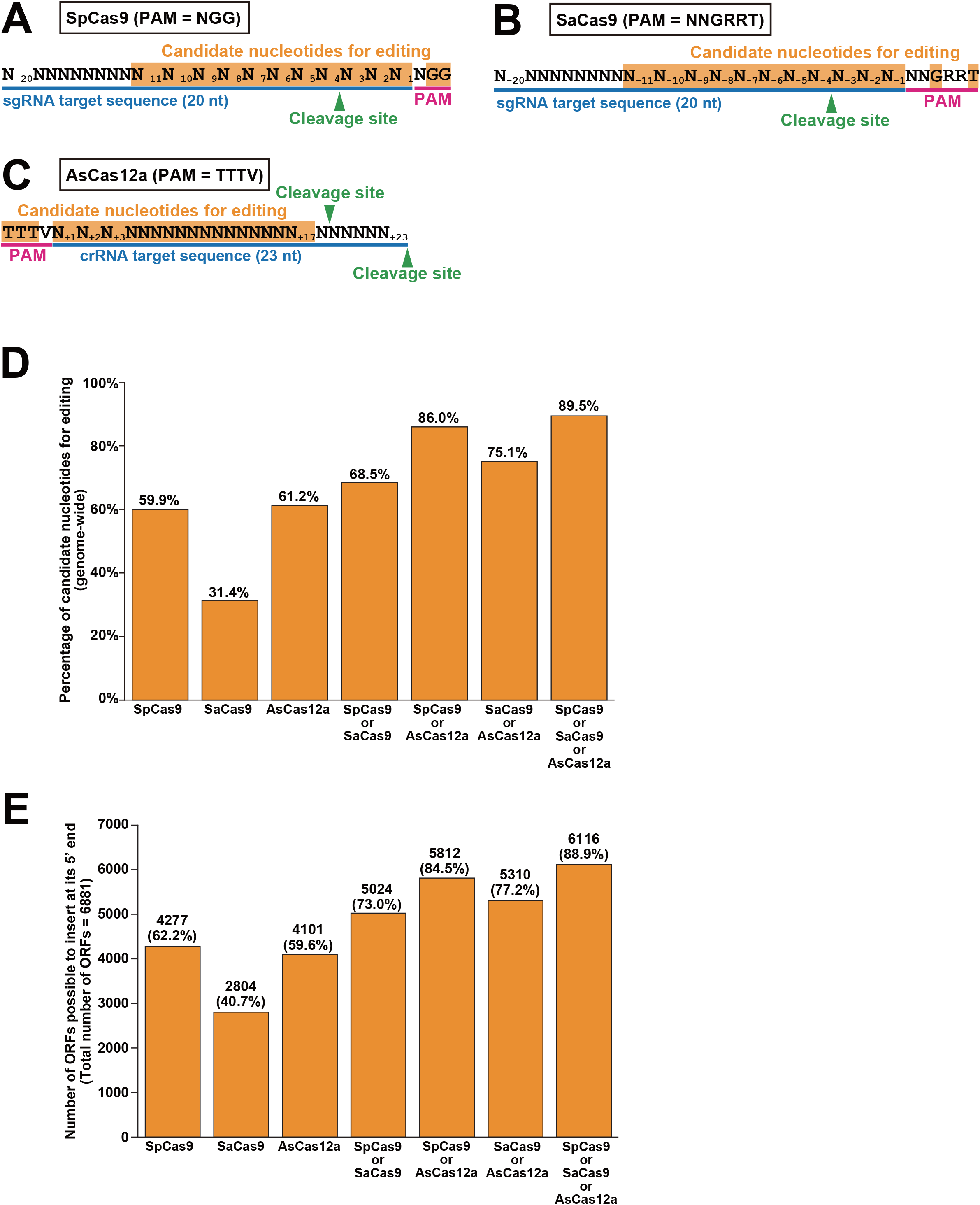
Estimation of editable nucleotides in the yeast genome and ORFs editable at their 5′ ends. (A, B, C) Definition of candidate nucleotides for editing. PAM sequence for each CRISPR/Cas system is underlined with magenta. The target sequence in sgRNA/crRNA is underlined with blue. Candidate nucleotides for editing with each CRISPR/Cas system are highlighted with orange. Green arrowheads indicate cleavage sites. (D) Fraction of nucleotides in the reference genome editable with each and all possible combinations of the three CRISPR/Cas systems. (E) Number of ORFs amenable to insertion at their 5′ ends with each and all possible combinations of the three CRISPR/Cas systems. Percentage of editable ORFs is shown in parentheses.

**Figure S2.**
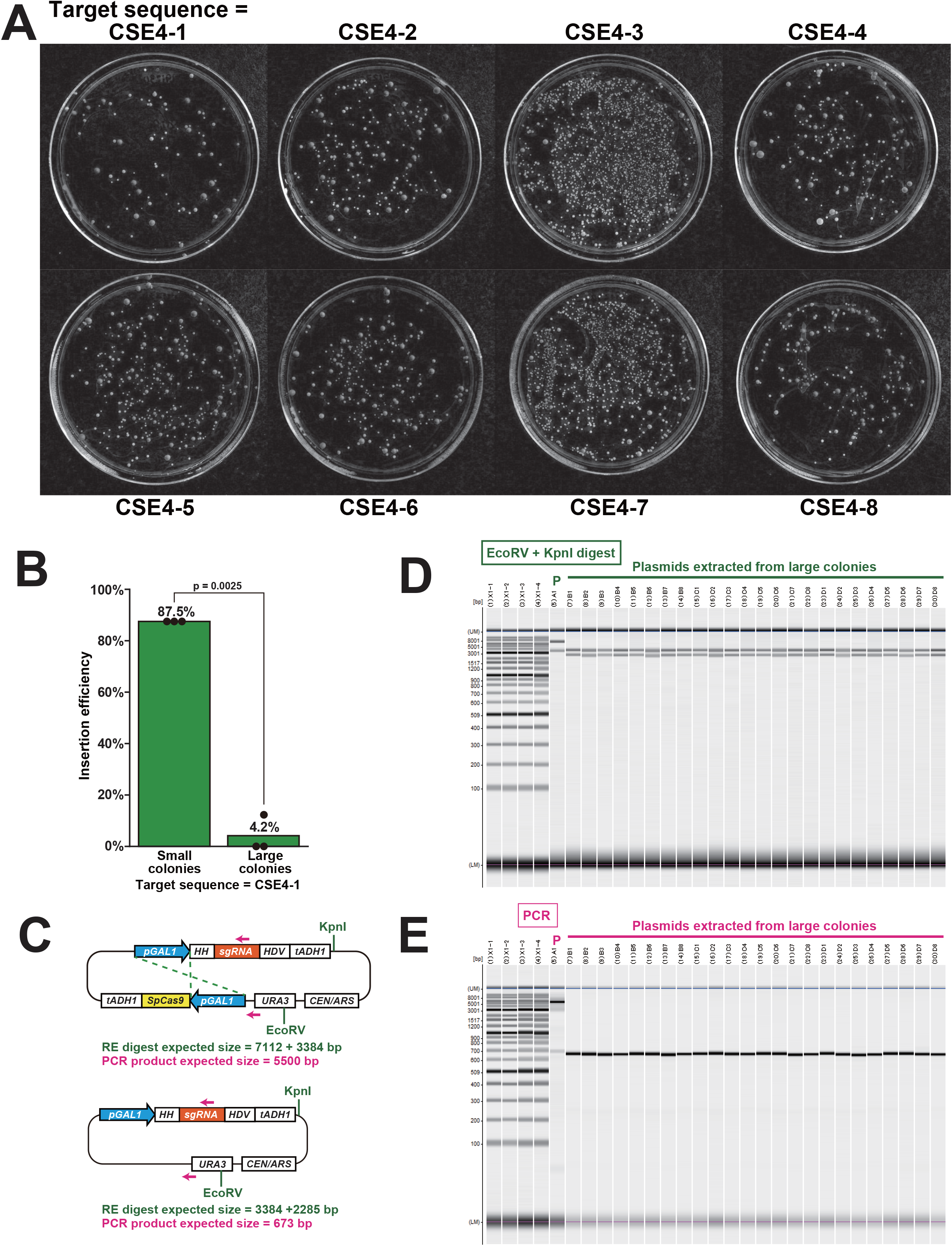
Characterization of large and small colonies obtained by transformation of genome-editing plasmids. (A) Representative images of the colonies on galactose-containing agar plates. The cells were transformed with the pGAL1-SpCas9 + pGAL1-sgRNA plasmid for insertion of mNeonGreen-encoding gene to *CSE4* (Figure 3). Each plate is labeled with the target sequence name shown in Figure 3A. (B) Insertion efficiency at the CSE4-1 target sequence. Green bars indicate the insertion efficiency (n = 8 for each of 3 biological replicates) of small colonies (left) and large colonies (right). The p-value of paired-samples t-test is shown. (C) Structures of the parental plasmid and its putative derivative generated via intramolecular recombination between the two *GAL1* promoters. Restriction enzyme sites used for plasmid DNA digestion are indicated in green. Magenta arrows indicate the positions of primers used for PCR. (D) Plasmid DNAs digested with EcoRV and KpnI analyzed by a microchip electrophoresis system. P, parental plasmid. (E) PCR products analyzed by a microchip electrophoresis system. P, parental plasmid.

**Figure S3.**
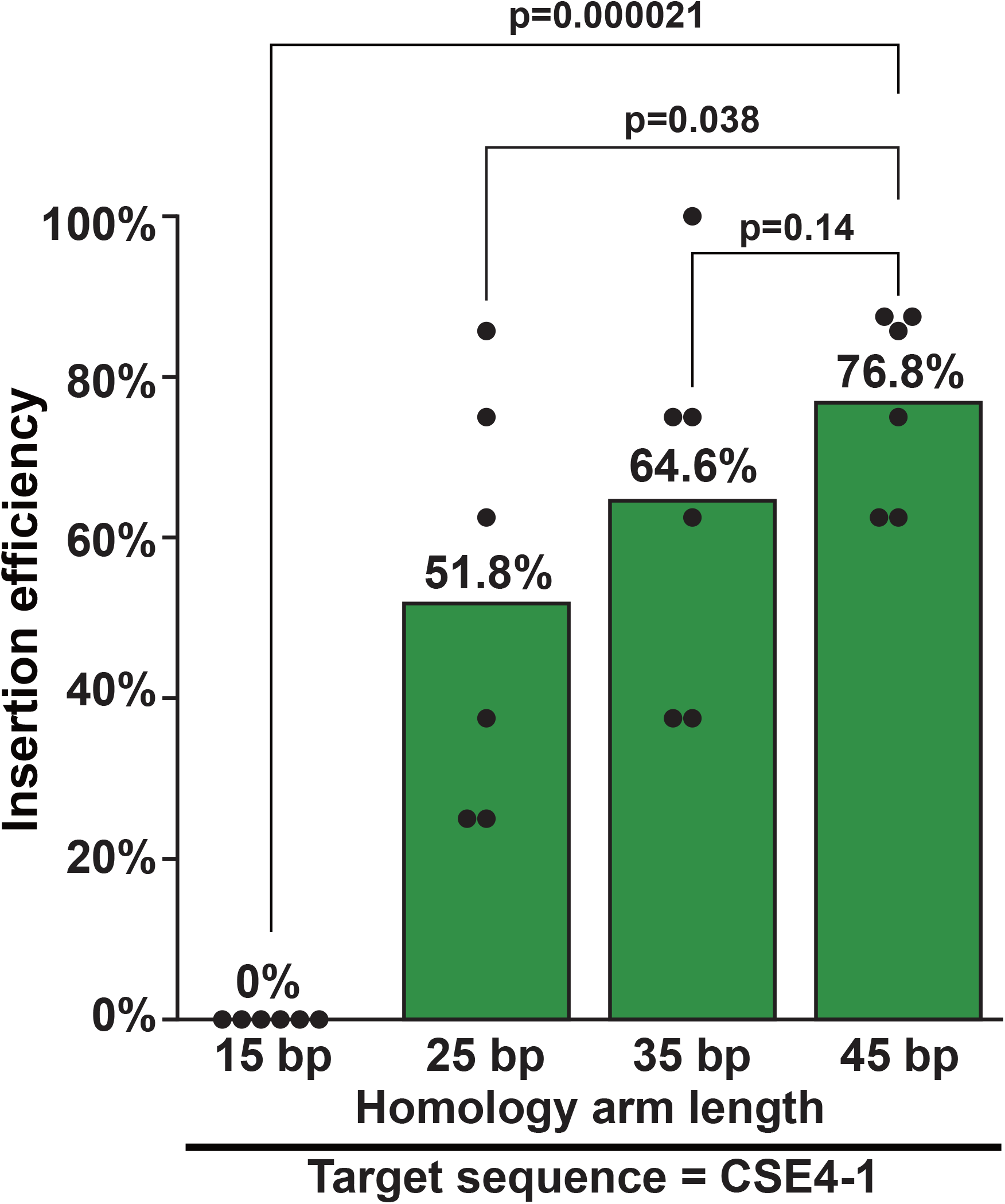
Effects of homology arm length on gene fragment insertion. Insertion efficiency at the CSE4-1 target sequence is indicated for donor PCR fragments harboring homology arms of four different lengths. Green bars indicate the insertion efficiency (n = 8 for each of 6 biological replicates). Only small colonies were examined.

**Figure S4.**
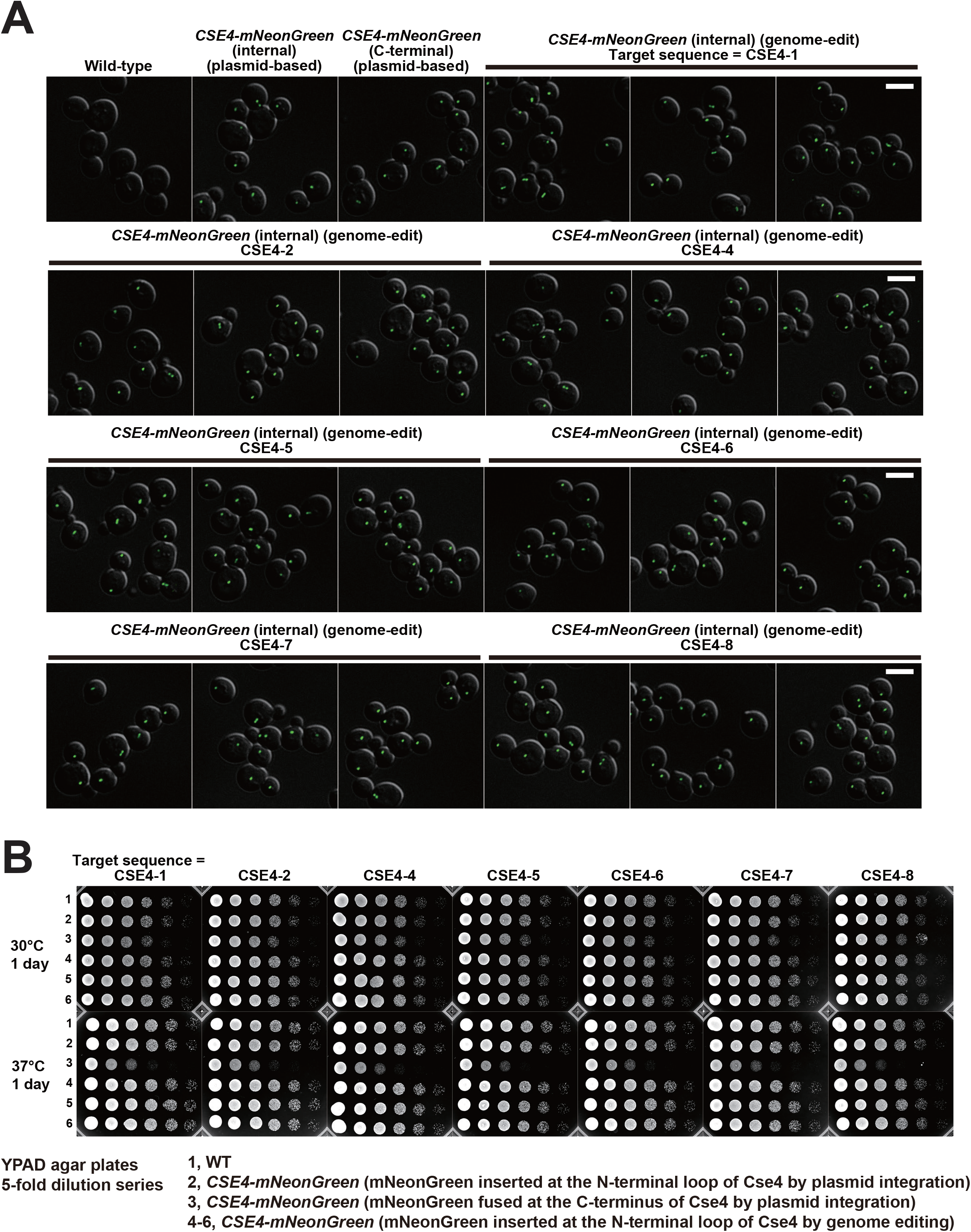
Characterization of *CSE4-mNeonGreen* cells generated by SpCas9-mediated genome editing. (A) Representative images of the wild-type cells, the *CSE4-mNeonGreen* cells generated by the conventional plasmid integration method, and the genome-edited *CSE4-mNeonGreen* cells. Images are composed by superimposition of DIC images (gray scale) and mNeonGreen fluorescent images (green). The target sequence names are shown above the images. Scale bar, 5 μm. (B) Images of cells grown on YPAD plates at 30°C and 37°C for a day. Overnight culture in YPAD liquid medium of each strain was diluted to the same cell density among the samples, serially diluted (5-fold), and spotted on YPAD agar plates (5 μL per spot).

**Figure S5.**
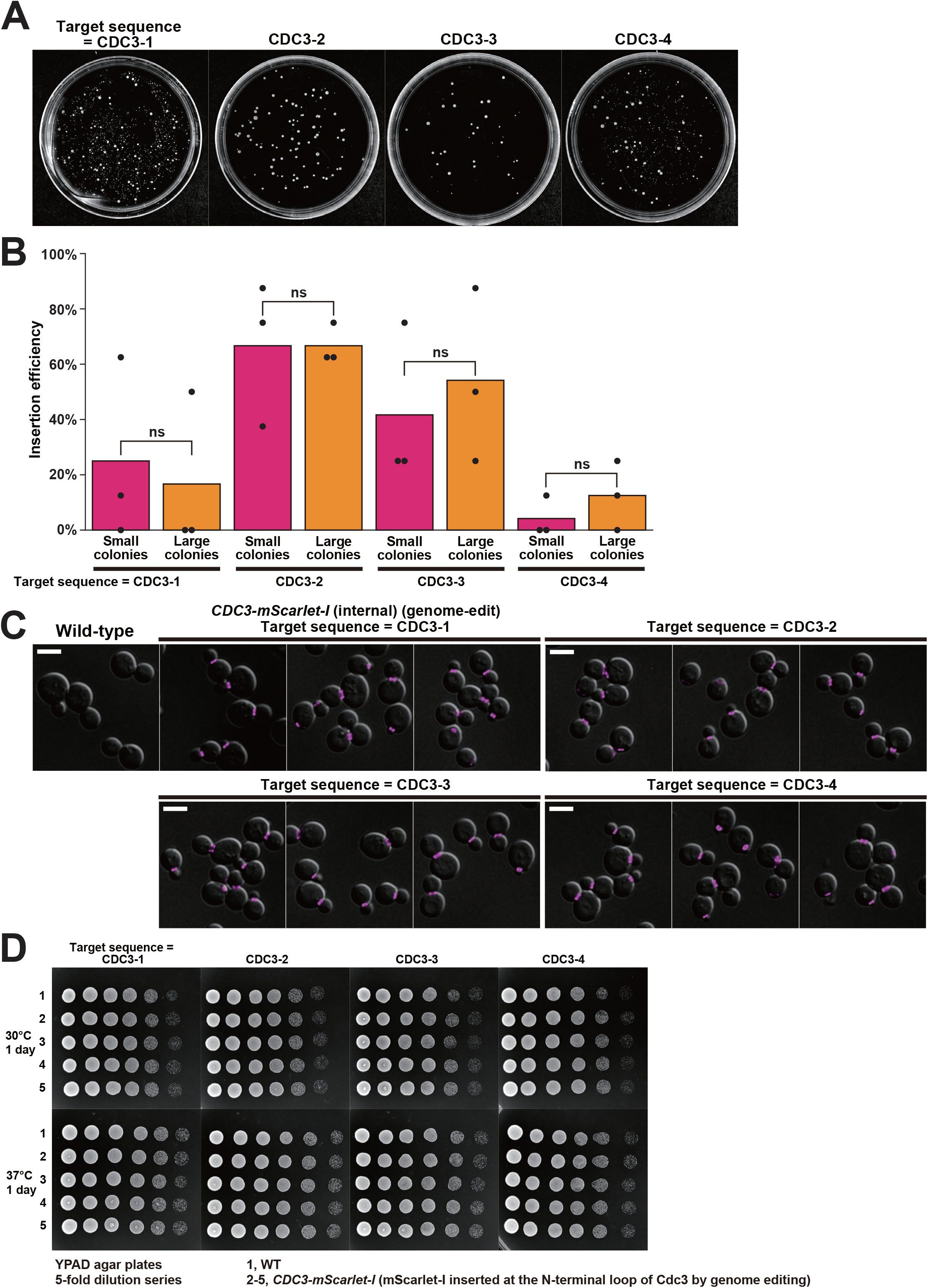
Characterization of *CDC3-mScarlet-I* cells generated by SpCas9-mediated genome editing. (A) Representative images of the colonies on galactose-containing agar plates. The cells were transformed with the pGAL1-SpCas9 + pSNR52-sgRNA plasmid. The target sequence names used for genome editing are shown above the plates. The positions of the target sequences are shown in Figure 4A. (B) Insertion efficiency at the 4 different target sequences in the *CDC3* gene. Magenta and orange bars indicate the insertion efficiency of small and large colonies, respectively (n = 8 for each of 3 biological replicates). For each target sequence, insertion efficiency does not show a statistically significant difference between small and large colonies (paired-samples t-test). (C) Representative images of the wild-type cells and the genome-edited *CDC3-mScarlet-I* cells. Images are composed by superimposition of DIC images (gray scale) and mScarlet-I fluorescent images (magenta). The target sequence names are shown above the images. Scale bar, 5 μm. (D) Images of cells grown on YPAD plates at 30°C and 37°C for a day. Overnight culture in YPAD liquid medium of each strain was diluted to the same cell density among the samples, serially diluted (5-fold), and spotted on YPAD agar plates (5 μL per spot).

**Figure S6.**
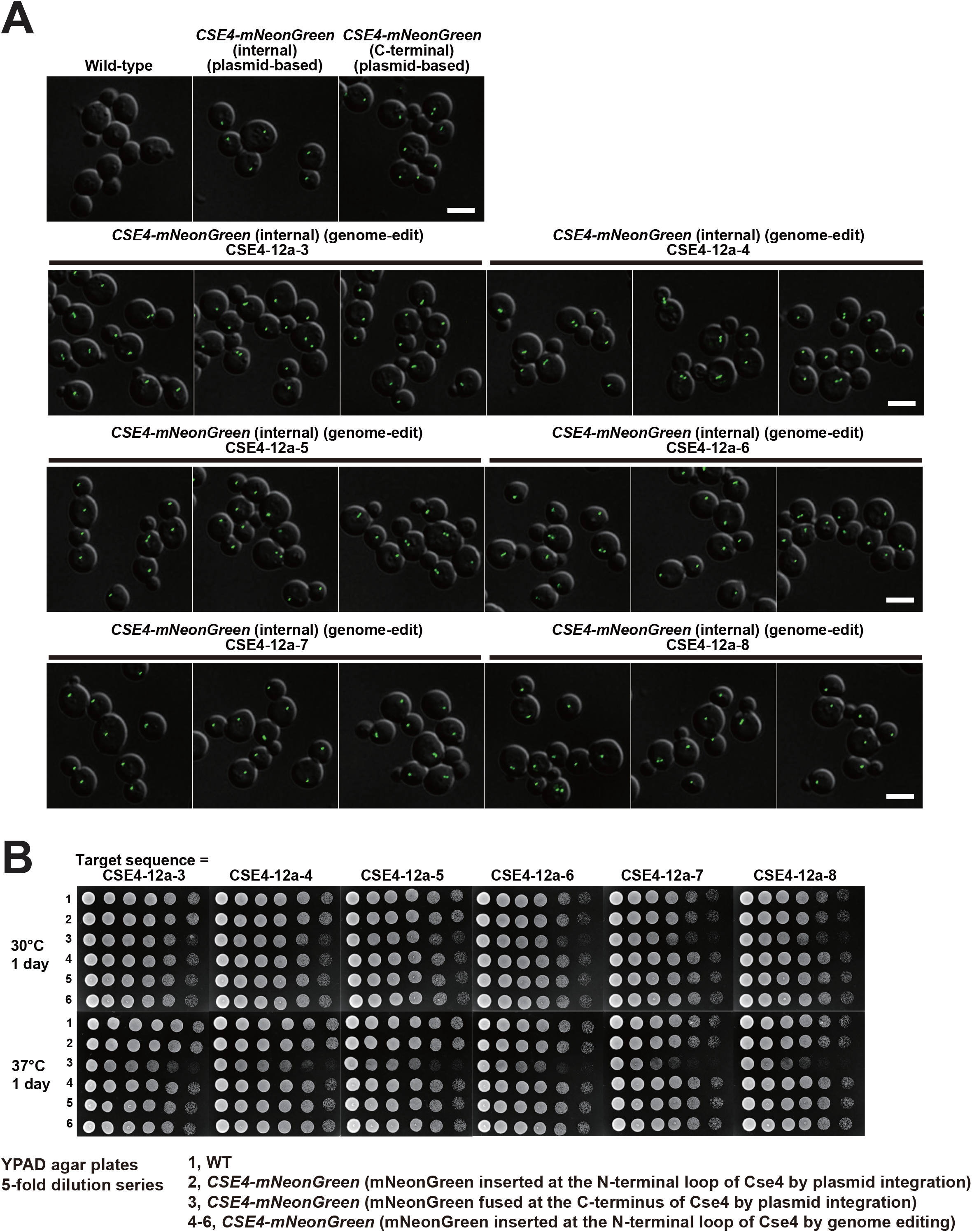
Characterization of *CSE4-mNeonGreen* cells generated by enAsCas12a-mediated genome editing. (A) Representative images of the wild-type cells, the *CSE4-mNeonGreen* cells generated by the conventional plasmid integration method, and the genome-edited *CSE4-mNeonGreen* cells. Images are composed by superimposition of DIC images (gray scale) and mNeonGreen fluorescent images (green). The target sequence names are shown above the images. Scale bar, 5 μm. (B) Images of cells grown on YPAD plates at 30°C and 37°C for a day. Overnight culture in YPAD liquid medium of each strain was diluted to the same cell density among the samples, serially diluted (5-fold), and spotted on YPAD agar plates (5 μL per spot).

**Figure S7.**
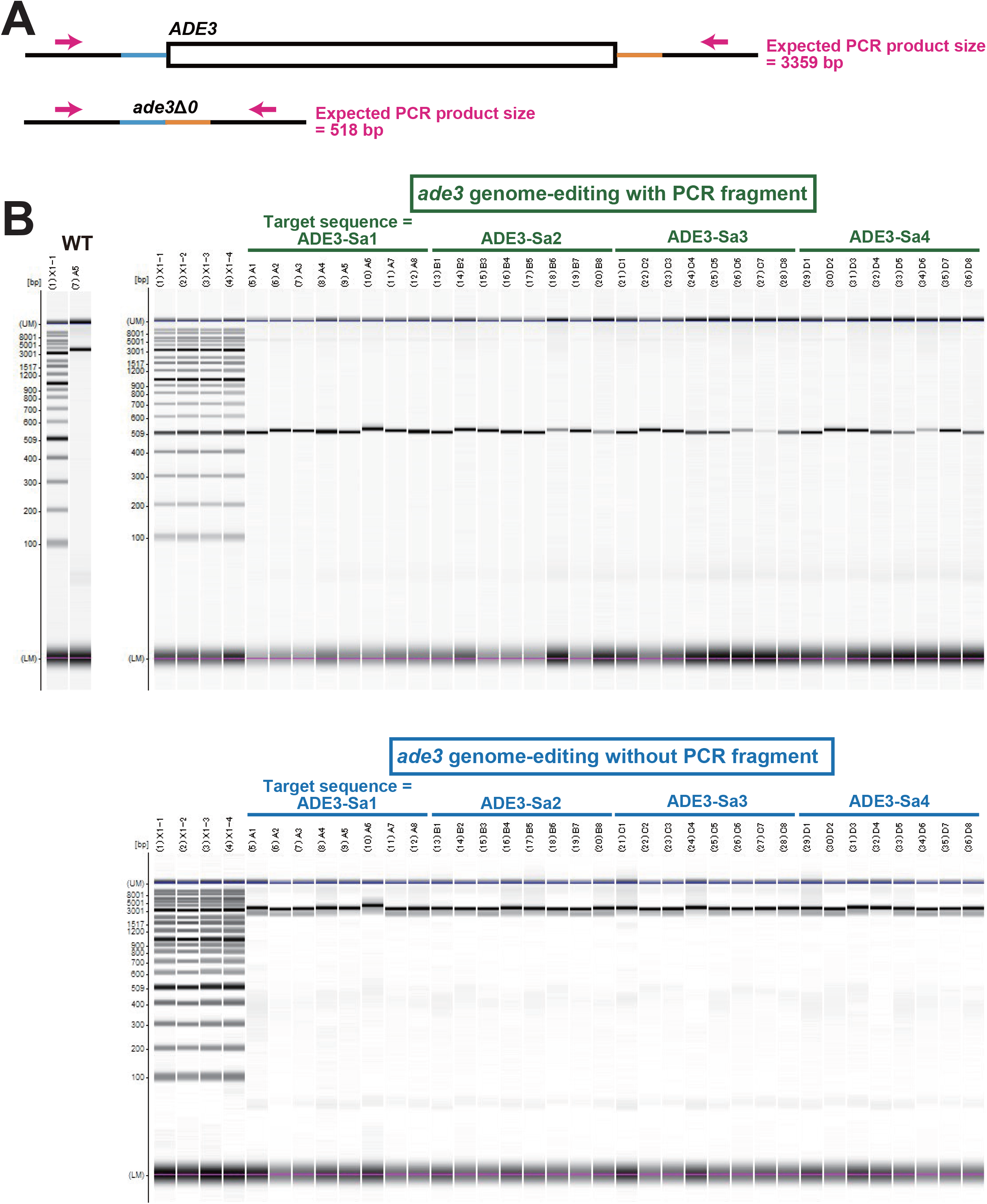
PCR characterization of white colonies obtained by SaCas9-mediated genome editing of *ADE3*. (A) Schematic representation of the *ADE3* locus (before genome editing) and the *ade3*Δ0 locus (after genome editing). The primer positions used for PCR check are indicated by magenta arrows. (B) PCR products from white colonies appeared on adenine-limited galactose-containing agar plates. Transformation was performed with (top) and without (bottom) the donor PCR fragment shown in Figure 6.

**Figure S8.**
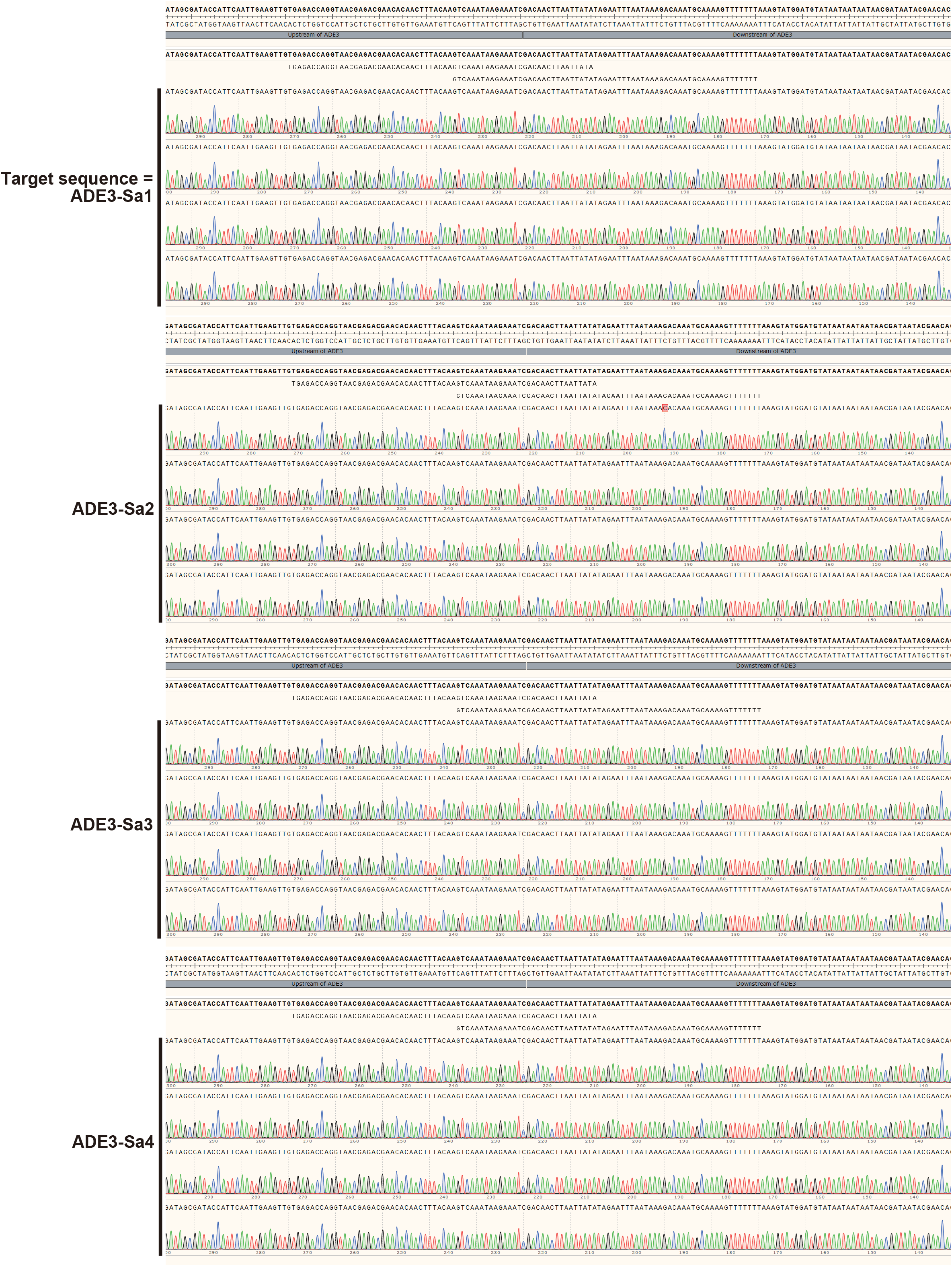
Nucleotide sequences of *ade3*Δ0 alleles generated by SaCas9-mediated genome editing with a donor PCR fragment. Nucleotide sequences are shown for the *ade3*Δ0 alleles generated by transformation of the individual genome-editing plasmids with the donor PCR fragment (Figure 6A). The expected *ade3*Δ0 sequence is shown at the top. Two horizontal gray bars are the upstream and downstream sequences of the *ADE3* gene.

**Figure S9.**
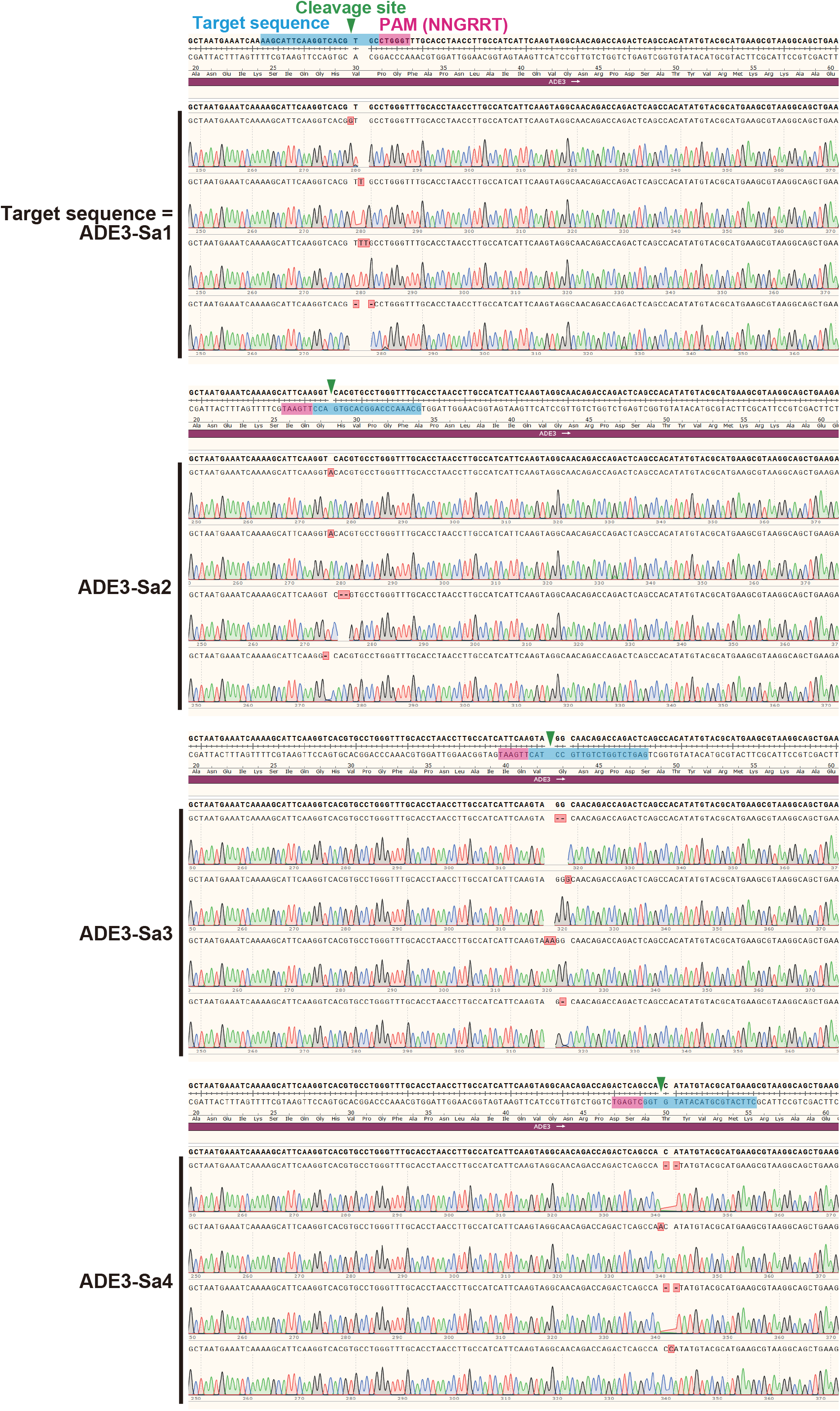
Nucleotide sequences of *ade3* alleles generated by genome editing without a donor PCR fragment. Nucleotide sequences are shown for the *ade3* alleles generated by transformation of the individual genome-editing plasmids without the donor PCR fragment (Figure 6A). At the top of each panel, the unedited *ADE3* sequence is shown. The target sequence and the PAM are highlighted with blue and magenta, respectively. Green triangles indicate the expected cleavage site by SaCas9. Inserted or deleted nucleotides are boxed by red and highlighted with pink in each sequence trace.

